# Quantifying multi-species microbial interactions in the larval zebrafish gut

**DOI:** 10.1101/2020.05.28.121855

**Authors:** Deepika Sundarraman, Edouard A. Hay, Dylan M. Martins, Drew S. Shields, Noah L. Pettinari, Raghuveer Parthasarathy

**Author notes:** These authors contributed equally to this work (Deepika Sundarraman, Edouard A. Hay, Dylan M. Martins). Department of Physics, University of Oregon, Eugene, OR 97403.

## Abstract

The microbial communities resident in animal intestines are composed of multiple species that together play important roles in host development, health and disease. Due to the complexity of these communities and the difficulty of characterizing them in situ, the determinants of microbial composition remain largely unknown. Further, it is unclear for many multi-species consortia whether their species-level makeup can be predicted based on an understanding of pairwise species interactions, or whether higher-order interactions are needed to explain emergent compositions. To address this, we examine commensal intestinal microbes in larval zebrafish, initially raised germ-free to allow introduction of controlled combinations of bacterial species. Using a dissection and plating assay, we demonstrate the construction of communities of one to five bacterial species and show that the outcomes from the two-species competitions fail to predict species abundances in more complex communities. With multiple species present, inter-bacterial interactions become weaker and more cooperative, suggesting that higher-order interactions in the vertebrate gut may stabilize complex communities.

## Introduction

Intestinal microbes exist in complex and heterogeneous communities of interacting, taxonomically diverse species. The composition of these communities varies across individuals and is crucial to the health of the host, having been shown in humans and other animals to be correlated with dietary fat uptake [1, 2], organ development [3, 4], immune regulation [5–10] and a wide range of diseases [11–20].

Despite the importance of intestinal communities, the determinants of their composition remain largely unknown. A growing number of studies map the effects of external perturbations, such as antibiotic drugs [21, 22] and dietary fiber [23] and fat [24, 25] on the relative abundance of gut microbial species. Intrinsic inter-microbial interactions, however, are especially challenging to measure and are important not only for shaping community composition in the absence of perturbations but also for propagating species-specific perturbations to the rest of the intestinal ecosystem.

The considerable majority of studies of the gut microbiota have been performed on naturally assembled microbiomes by sequencing DNA extracted from fecal samples, an approach that provides information about the microbial species and genes present in the gut, but that imposes several limitations on the inference of inter-species interactions. The high diversity of natural intestinal communities, and therefore the low abundance of any given species among the multitude of its fellow residents, implies that stochastic fluctuations in each species’ abundance will be large, easily masking true biological interactions. The accuracy of inference is considerably worse if only relative, rather than absolute, abundance data is available [26–29], as is typically the case in sequencing-based studies. Finally, we note that fecal sampling assesses only the microbes that have exited the host, which may not be representative of the intestinal community [30].

An alternative approach to using DNA sequencing and naturally assembled host-microbiota systems is to build such systems from the bottom-up using model organisms. This is accomplished by using techniques for generating initially germ-free animals, and well-defined sets of small numbers of microbial species, and then measuring the populations of these species resident in the intestine. Recent work along these lines has been performed using the nematode *Caenorhabditis elegans* [31] and the fruit fly *Drosophila melanogaster* [32, 33]. However, as described further below, these studies imply different principles at play in the different systems. Moreover, it is unclear whether conclusions from either model platform translate to a vertebrate gut, which has both greater anatomical complexity and more specific microbial selection [34]. To address this, we measure bacterial interactions in larval zebrafish (Fig 1A), a model vertebrate organism amenable to gnotobiotic techniques [35–38], which has enabled in earlier work that investigated pairs of bacterial species the discovery of specific interbacterial competition mechanisms related to intestinal transport [39, 40]. The experiments described below involve several hundred fish, each with 1-5 resident bacterial species, enabling robust inference of inter-species interactions.

**Fig 1.**
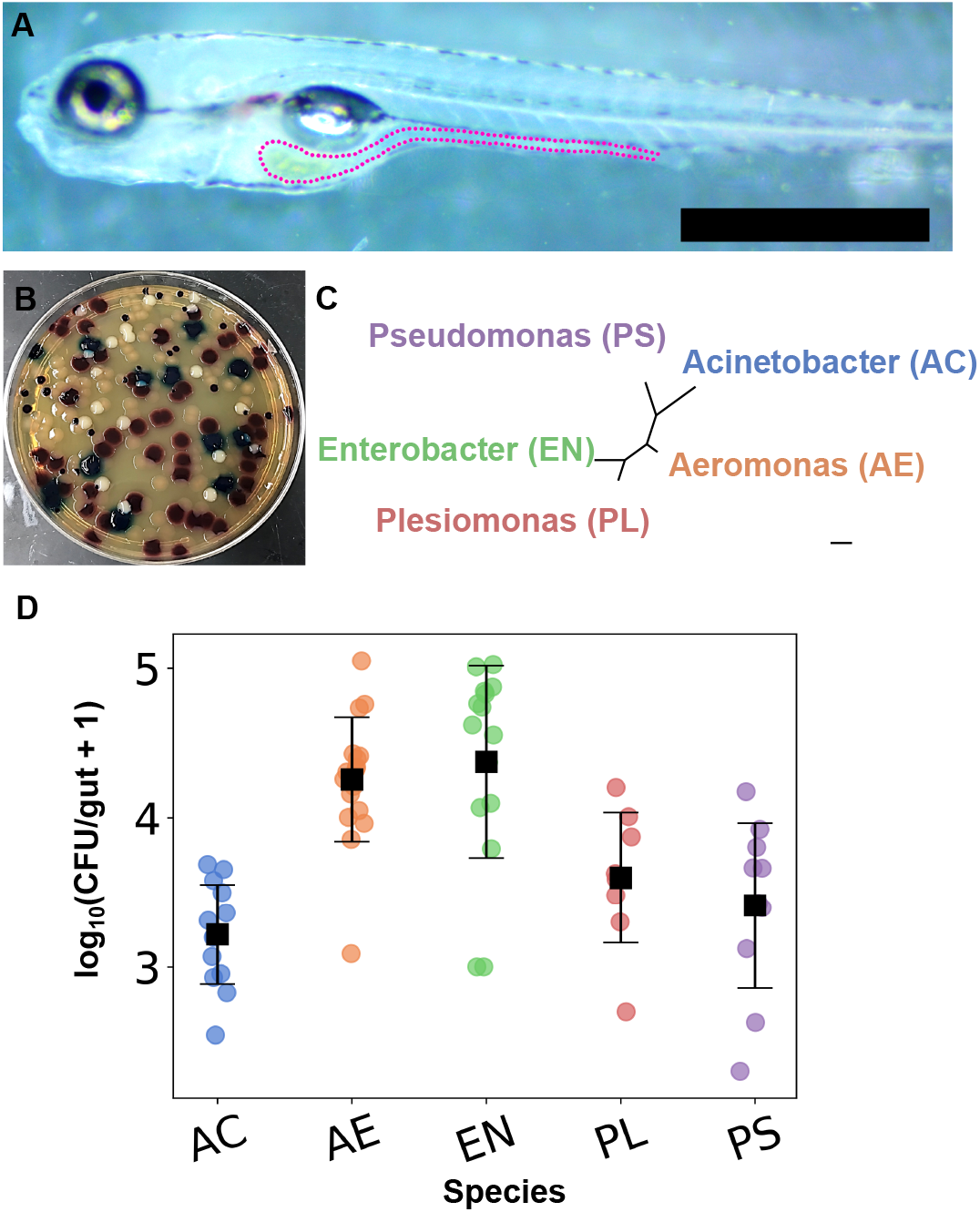
The five chosen commensal species are robust colonizers of the larval zebrafish intestine. (A) A 7 day post fertilization larval zebrafish, with a dotted line outlining the intestine. Scale bar: 500 μm. (B) Chromogenic agar plate showing colonies of all the candidate five species (AC: milky opaque, AE: reddish purple, EN: blue, PL: dark purple and PS: colorless translucent) (C) A phylogenetic tree of the five bacterial species examined in this study, calculated from 16S rRNA sequences using PhyML. Scale bar: 0.05 nucleotide substitutions per site. (D) The abundance per zebrafish gut of each of the five bacterial species when colonized in mono-association with the host, assessed as colony forming units (CFU) from plated gut contents. Each circular datapoint is a CFU value from an individual fish (N = 13, 17, 15, 8, and 10, from left to right), with the mean and standard deviation indicated by the square markers and error bars.

The ability to quantify species abundance and to manipulate it by controlled addition or subtraction of species is commonplace in macroscopic ecological investigations. Its implementation here enables connections between intestinal microbiome research and a large literature on ecosystem dynamics. An issue whose importance has been realized for decades is the extent to which interspecies interactions are pairwise additive, or whether higher-order (often called indirect) interactions are necessary to explain community structure [41, 42]. Pairwise additivity, if dominant, simplifies the prediction of ecosystem composition, which would be desirable for therapeutic applications of microbiome engineering. Higher order interactions may stabilize multi-species communities according to several recent theoretical models described further in the Discussion [43–46], implying that quantifying and controlling indirect effects may be necessary for reshaping gut microbiomes.

Whether host-associated or not, microbial communities have shown a variety of interaction types. A classic study involving cultured protozoan species found good agreement between the dynamics of four-species consortia and predictions derived from measurements of pairs of species [47]. Similarly, Friedman and colleagues showed that the outcomes of competitions among three species communities of soil-derived bacteria could be simply predicted from the outcomes of pairwise combinations [48]. In contrast, experiments based on the cheese rind microbiome found significant differences in the genes required for a non-native *E. coli* species to persist in a multi-species bacterial community compared to predictions from pairwise coexistence with community members [49]. A closed ecosystem consisting of one species each of algae, bacteria, and ciliates exhibited a strong non-pairwise interaction, in which the bacterium is abundant in the presence of each of the algae or ciliate alone, but is subject to strong predation in the three-species system [50].

Within animals, the interaction types observed in the few studies to date that make use of controlled microbial communities in gnotobiotic hosts are also disparate. Competitive outcomes of three-species communities from subsets of eleven different bacterial species in the gut of the nematode *C. elegans* could be predicted from the outcomes of two-species experiments, with indirect effects found to be weak [31]. In contrast, work using well-defined bacterial assemblies of up to five species in the fruit fly *D. melanogaster* found strong higher order interactions governing microbe-dependent effects on host traits such as lifespan [32].

To our knowledge, there have been no quantitative assessments of inter-bacterial interactions using controlled combinations of microbial species in a vertebrate host, leaving open the question of whether higher order interactions are strong, or whether pairwise characterizations suffice to predict intestinal community structure. We therefore examined larval zebrafish, inoculating initially germ-free animals with specific subsets of five different species of zebrafish-derived bacteria and assessing their subsequent absolute abundances. Though the number of species is considerably fewer than the hundreds that may be present in a normal zebrafish intestine, it is large enough to sample a range of higher-order interactions, yet small enough that the number of permutations of species is tractable.

As detailed below, we find strong pairwise interactions between certain bacterial species. However, we find weaker interactions and a greater than expected level of coexistence in fish colonized by four or five bacterial species. This suggests that measurements of pairwise inter-microbial interactions are insufficient to predict the composition of multispecies gut communities, and that higher-order interactions may dampen strong competition and facilitate diversity in a vertebrate intestine.

## Materials and methods

### Animal Care

All experiments with zebrafish were done in accordance with protocols approved by the University of Oregon Institutional Animal Care and Use Committee and following standard protocols [51].

### Gnotobiology

Wild-type (ABCxTU strain) larval zebrafish (*Danio rerio*) were derived germ-free as described in [36]. In brief, embryos were washed at approximately 7 hours post-fertilization with antibiotic, bleach, and iodine solutions and then moved to tissue culture flasks of 15mL sterile embryo medium solution with approximately 1mL of sterile solution per larva. The flasks were then stored in a temperature-controlled room maintained at 28° C.

### Bacterial Strains and Culture

The five bacterial strains used in this study, namely *Aeromonas sp*. (ZOR0001), *Pseudomonas mendocina* (ZWU0006), *Acinetobacter calcoaceticus* (ZOR0008), *Enterobacter sp*. (ZOR0014), and *Plesiomonas sp*. (ZOR0011) were originally isolated from the zebrafish intestine and have been fluorescently labelled to express GFP and dTomato facilitating their identification in our experimental assays [52, 53]. Stocks of bacteria were maintained in 25% glycerol at –80° C.

### Inoculation of tissue culture flasks

One day prior to inoculation of the tissue culture flasks, bacteria from frozen glycerol stocks were shaken overnight in Lysogeny Broth (LB media, 10 g/L NaCl, 5 g/L yeast extract, 12 g/L tryptone, 1 g/L glucose) and grown for 16 h overnight at 30°C. Samples of 1mL of each of the overnight cultures were washed twice by centrifuging at 7000g/rpm for 2 min, removing the supernatant, and adding 1mL of fresh sterile embryo media. At 5 dpf, the tissue culture flasks were inoculated with this solution at a concentration of 10^6^ CFU/mL. For each of the competition experiments involving 2, 4 and 5 bacterial species, equal concentrations were inoculated into the flasks. After inoculation, the flasks were maintained at 30°C until dissection at 7 dpf.

### Dissection and Plating

To determine the intestinal abundance of bacterial species, dissections of larval zebrafish were performed at 7 dpf. Zebrafish were euthanized by hypothermal shock. Intestines were removed by dissection and placed in 500μL of sterile embryo media and homogenized with zirconium oxide beads using a bullet blender. The homogenized gut solution was diluted to 10^−1^ and 10^−2^, and 100μL of these dilutions were spread onto agar plates. Flask water was diluted to 10^−4^, and 100μL of these dilutions were spread onto agar plates. For mono- and di-associated inoculations, tryptic soy agar (TSA) plates were used in which fluorescence could be used to differentiate up to two species. For inoculations of more than two species, Universal HiChrome Agar (Sigma-Aldrich) plates were used, allowing for visual differentiation of each species using a colorimetric indicator. The abundances of each of the species in the zebrafish gut was determined by counting the colony forming units on the plates. These abundances for different experiments are provided in S1 File.

### In vitro competition experiments

To determine the in vitro competition coefficients, all the different pairwise combinations of the five species were grown in overnight cultures of LB media as above. On the following day, cultures were plated at 10^−7^ or 10^−6^ dilutions, depending on the ability to detect both species in a given dilution. Abundances were obtained by counting the number of CFUs of each species on the plates. These are provided in S2 File.

## Results

Zebrafish (Fig 1A) were derived to be germ-free, and then were inoculated at 5 days post-fertilization (dpf) with the desired combination of microbial species by addition of bacteria to the flasks housing the fish. Approximately 48 hours later, fish were euthanized and their intestines were removed by dissection. Intestines and their contents were homogenized, diluted, and plated onto chromogenic agar (Methods). Secreted enzymes from each of the five candidate bacterial species generate particular colors due to substrates in the chromogenic medium, allowing quantification of colony forming units (CFUs) and therefore absolute intestinal abundance (Fig 1B). All abundance data are provided in S1 File.

The five species examined were selected as diverse representatives of genera commonly found in the zebrafish intestine. Full names and species identifiers are given in Methods; we will refer to these through most of the text by genus name or two letter abbreviation: *Acinetobacter* (AC), *Aeromonas* (AE), *Enterobacter* (EN), *Plesiomonas* (PL), and *Pseudomonas* (PS) (Fig 1C). As expected given their association with the zebrafish gut microbiome, each species in mono-association, i.e. as the sole species inoculated in germ-free fish, colonizes robustly to an abundance of 10^3^-10^4^ CFU/gut, corresponding to an in vivo density of approximately 10^9^-10^10^ bacteria/ml (Fig 1D).

### Pairwise Interactions in Di-associations

We first examined all ten possible co-inoculations of two species, which enables assessment of pairwise interactions in the absence of higher-order effects. Intestinal CFU data shows a wide range of outcomes for different species pairs. As exemplars, the CFUs per gut for each of two species, AC and EN, in the presence of each of the other four are displayed in Figs 1A and B respectively. The abundance of AC is similar in the presence of any second species to its value in mono-association. In contrast, the mean EN abundance is similar to its mono-association value if co-inoculated with PL or PS, about 10 times lower if co-inoculated with AC, and over two orders of magnitude lower if co-inoculated with AE, implying in the latter cases strong negative interactions.

Parameterizing the strength of interactions between species is necessarily model dependent, contingent on the functional form of the relationship between one species’ abundance and the other’s. We show that the conclusions we reach regarding interaction strengths, especially their shifts when multiple species are present, are qualitatively similar and therefore robust for a wide range of models. We first consider a phenomenological interaction coefficient 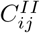 that is linear in log-abundance, characterizing the effect of species j on species i as:

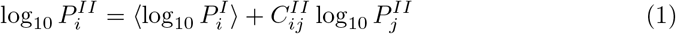

where *P_i_* denotes the abundance of species i and the superscript I or II denotes a mono- or di-association experiment. This form is motivated by the distribution of gut bacterial abundances being roughly log-normal, with species addition capable of inducing orders-of-magnitude changes (Figs 2A,B). This 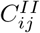 can be derived as the interaction parameter in a competitive Lotka-Volterra model modified to act on log-abundances (see S1 Text). Qualitatively, a positive *C_ij_* implies that the abundance of species i increases in the presence of j. Similarly a negative *C_ij_* indicates that the abundance of species i declines in the presence of species j. Subsampling from the measured sets of bacterial abundances gives the mean and standard deviation of the estimated interaction parameters (see S1 Text).

**Fig 2.**
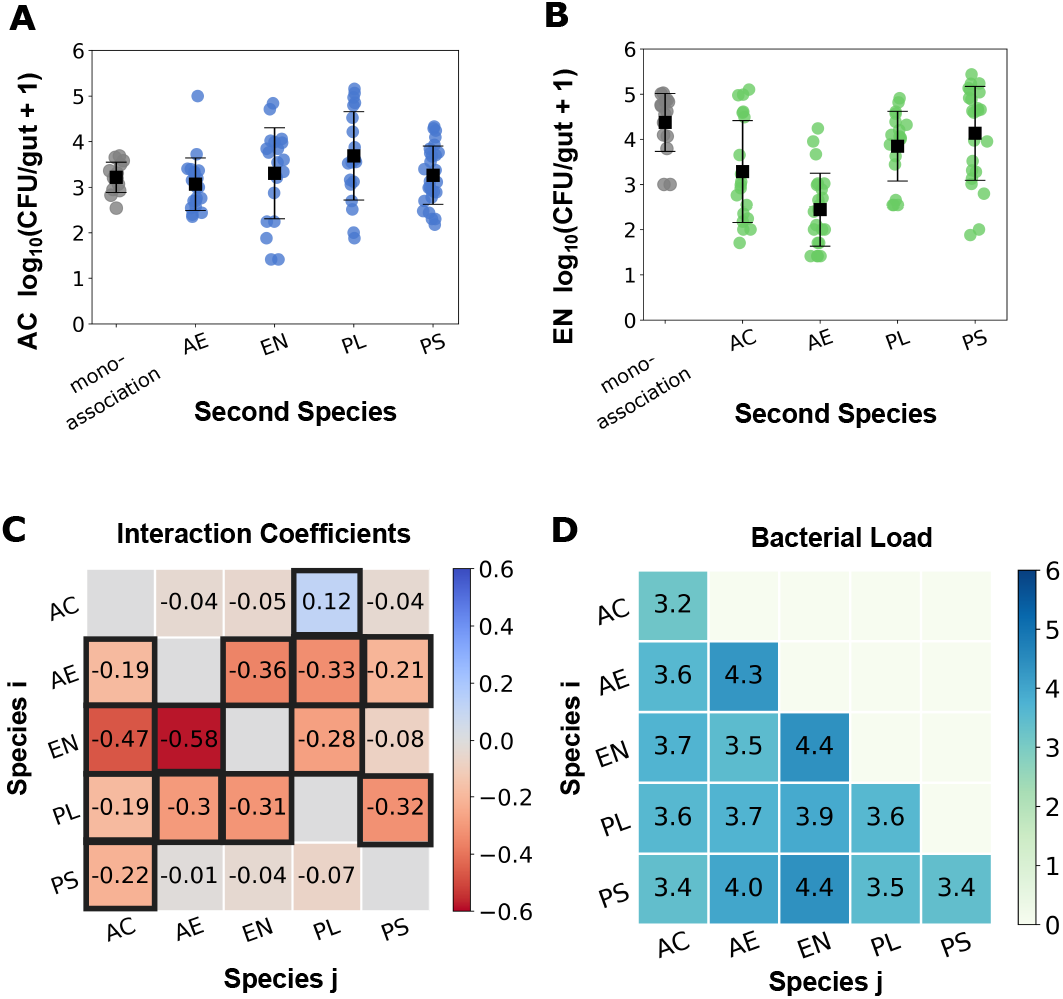
Strong negative pairwise interactions dominate di-association experiments. Abundances per zebrafish gut of (A) AC and (B) EN in mono-association (grey) and in di-association with each of the other bacterial species (blue/green). Each circular datapoint is a CFU value from an individual fish ((A) N = 13, 21, 19, 20, and 27 and (B) N = 15, 19, 22, 18, and 23 from left to right), with the mean and standard deviation indicated by the square markers and error bars. (C) Matrix of pairwise interaction coefficients 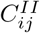 characterizing the effect of species j on the abundance of species i. Coefficients that differ from zero by more than three standard deviations (provided in S2 Fig) are outlined in black. (D) The average bacterial load per zebrafish in each of the di-association combinations, expressed as log_10_ of total CFUs. The standard deviations are between 0.3 and 1.1 and are displayed in S1 Text. Values on the diagonal are the mono-association load for each of the five species.

We plot in (Fig 2C) the 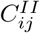 defined by Eq. 1 calculated from all di-association data of all species pairs (N = 190 fish in total). For determining 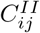, we only use data from fish in which both species were detected so that abundance changes of one species can definitively be ascribed to the presence of the other within the gut. Uncertainties in 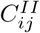 are estimated from bootstrap subsampling (see S1 Text). The interactions are predominantly negative, with thirteen out of twenty coefficients differing from zero by over three standard deviations. The total bacterial load, i.e. the sum of the bacterial abundances, is similar for all the di-associations suggesting that the interaction effects do not stem from changes in intestinal capacity (Fig 2D).

Though the physical and chemical environment of the zebrafish gut is likely very dissimilar to test tubes of standard growth media, we examined abundances of each of the pairs of species in in vitro competition experiments, growing overnight cultures in Lysogeny broth (LB) media and plating for CFUs (see Methods). Assessing 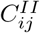 as above, we find, as expected, that interaction coefficients calculated from the in vitro experiments are markedly different than those measured in vivo (see S3 Fig) and (Fig 3B). Our characterizations of interactions within the zebrafish gut are not qualitatively altered by using a more general power law model to compute 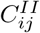 from absolute abundance data, discussed below (Interactions under more general models) following the presentation of measurements of interactions between more than two species.

**Fig 3.**
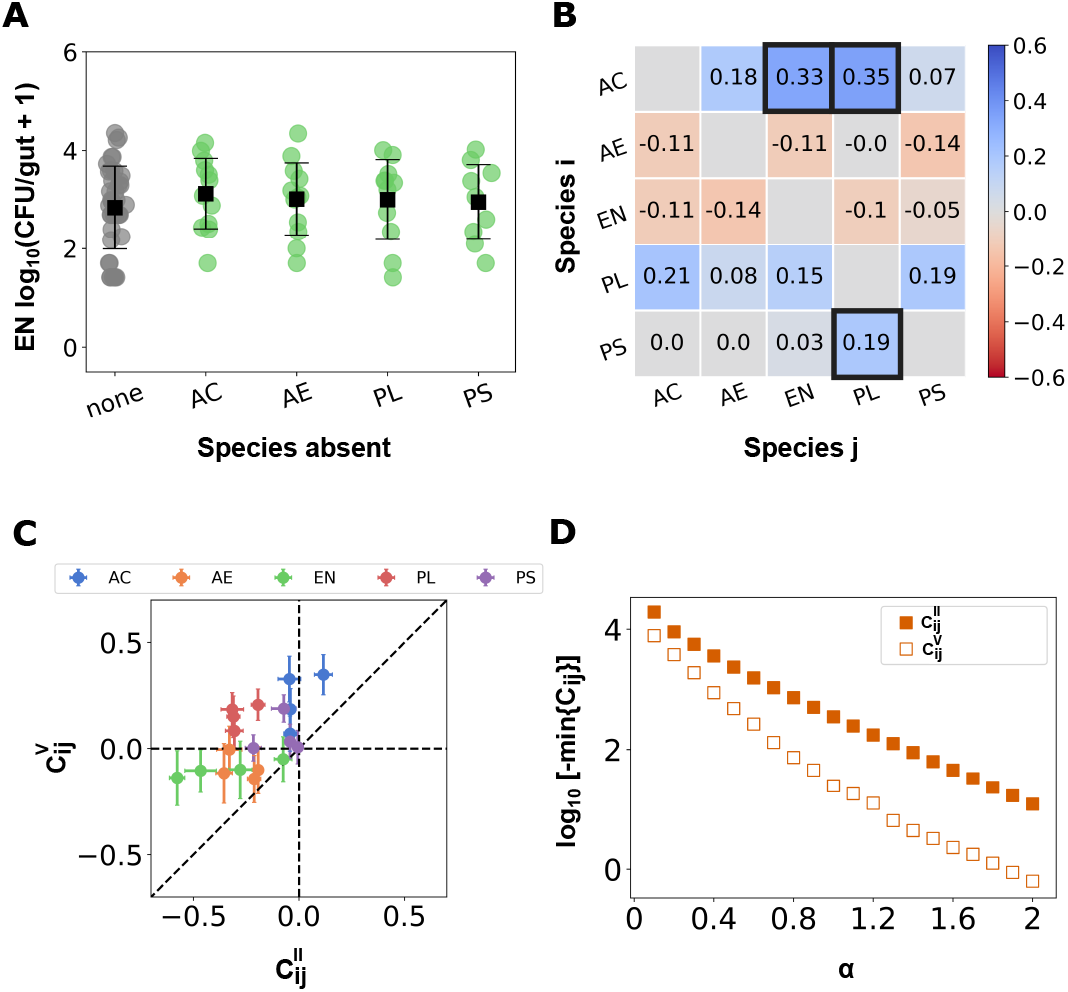
Weak pairwise interactions in five-species experiments. (A) Abundance per zebrafish gut of one of the bacterial species, EN, when all five species are co-inoculated (gray) and in each four species co-inoculation experiment (green) with the omitted species indicated on the axis. Each circular datapoint is a CFU value from an individual fish (N = 40, 12, 12, 11, and 9, from left to right), with the mean and standard deviation indicated by the square marker and error bars. (B) Matrix of pairwise interaction coefficients 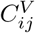 when 5 bacterial species are present. The coefficients outlined in black differ from zero by over three standard deviations (see S4 Fig). (C) The pairwise interaction coefficients inferred from 4-5 species experiments versus those from 1-2 species experiments. The colors label species i for each interaction pair. (D) The minimum interaction coefficient calculated from a power-law interaction model for different values of the exponent *α* for the 1-2 species (square filled markers) and the 4-5 species (square markers) experiments.

### Pairwise Interactions in Multi-species Communities

To assess whether the strong competitive interactions we found in two-species experiments are conserved in multi-species communities, we quantified pairwise interactions in experiments inoculating fish with four or five bacterial species. To assess 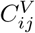, we adopted a method similar to the leave-one-out approach often used in macroscopic ecological studies, dating at least to classic experiments in which single species were removed from tide pools and the abundances of the remaining species were measured to evaluate inter-species interactions [54]. Here, we performed co-inoculation experiments leaving out one of the five species of bacteria and compared intestinal abundances for these four-species communities to those measured in five-species co-inoculation experiments.

In approximately N = 10 fish each, we performed all five different co-inoculations of four bacterial species. The difference in the abundance of species i in fish inoculated with all five species compared to fish inoculated with four, missing species j, gives a measure of the impact of species j on species i in the multi-species environment. As an example, EN abundance in inoculations lacking AC, AE, PL, and PS, and in five-species inoculations, are shown in Fig 3A. In contrast to the di-association experiments (Fig 2B), we see that EN does not show large abundance differences, in either its mean or its distribution, as a result of any fifth species being present.

Again, a variety of options are possible for quantifying interaction coefficients in the multi-species system. We first consider interaction coefficients as modifying mean-log-abundances, analogous to the pairwise model of Eq. 1:

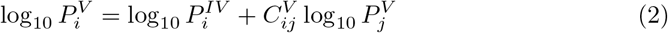

The interaction coefficients 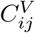 that we obtain, displayed in Fig 3B, are different and in general considerably weaker than those found in the two-species case Fig 2C. There are only three interactions that differ from zero by over three standard deviations. Strikingly, all three of these interactions are positive. This shift towards weaker and more positive interactions between the two-species and multi-species interactions is further illustrated in Fig 3C in which the multi-species interaction coefficients, 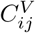 are plotted against the 2-species interaction coefficients, 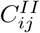.

### Interactions under more general models

As noted, a model that is linearly additive in logarithmic abundances is only one of an infinite number of choices, and moreover may not adequately capture the complexity of interactions in the gut. Earlier experiments investigating the spatial structure of specific microbial communities in the larval zebrafish intestine have shown that species such as AE, EN and PS form dense three-dimensional aggregates [55]. The size and location of aggregates and the locations of cells, conspecific or otherwise, within these aggregates may impact their interactions in ways that could be sub-linear, linear, or super-linear in population size. Previous work has also established that gut bacteria may also influence intestinal mechanics [40], highlighting one of many possible indirect interaction mechanisms whose functional forms are unknown. Furthermore, other studies have shown that different modes of physical and chemical communication could result in long range interactions between different species [56–58]. To address these possibilities, we evaluated species interactions with a more general power law model, wherein the interaction effects between species could be non-linear in the abundance of the effector species. Here the interaction coefficient *C_ij_* depends on a power, *α*, of the abundance of the effector species j, which we evaluate in the range *α* = 0.1 to 2, spanning sub-linear and super-linear interactions. Modified versions of Eq. 1 and 2 give:

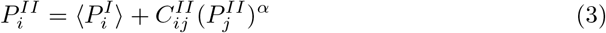

and

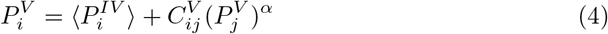

from which we can evaluate 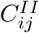 and 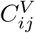 respectively. Note that *α* = 1 in Eq. 3, 4, i.e. interactions that are linear in abundance, is simply the steady-state behavior of the competitive Lotka Volterra model commonly used in population modeling and are shown in S6 Fig. We provide the 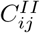 and 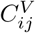 for several different α in S5 Fig. Throughout, as in the logarithmic model shown above, pairwise interactions in di-association are in many cases strongly negative, while the multi-species interactions are weaker. This is summarized by studying the trends in the most negative 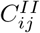 and 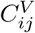 for different values of *α*, depicted in Fig 3D, which shows that for all *α* the strongest 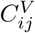 is significantly weaker than the strongest 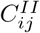, suggesting that our results are robust to choice of model.

### Five-species Coexistence

We next consider co-inoculation of all five bacterial species. Examination of over 200 fish shows a large variety in abundances, depicted in Fig 4A as the relative abundance of each species in each larval gut. Multiple species are able to coexist, with the median number of species present being 4 (Fig 4B). The mean total bacterial load as well as its distribution (Fig 4C) is similar to the mean and distribution of the mono- and di-association experiments, as well as four-species co-inoculation experiments discussed earlier. We calculated the expected abundance of each bacterial species, if the interactions governing the five-species community were simply a linear combination of the pairwise interactions governing di-associations, 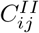. Any of the additive models we evaluated can be extended to combinations of species. Considering the model focused on above, with interaction coefficients modifying log abundances, the predicted abundance of species i in the presence of another species j is given by

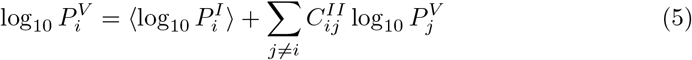

where the superscript V denotes the five-species co-inoculation experiment. A model linear in species abundance (*α* = 1) is also considered in the S1 Text), and gives qualitatively similar outputs and conclusions. Sampling from the measured distributions of each of the interaction coefficients and the mean abundance in mono-association allows calculation of the distribution of expected 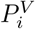 values (see S5 File).

**Fig 4.**
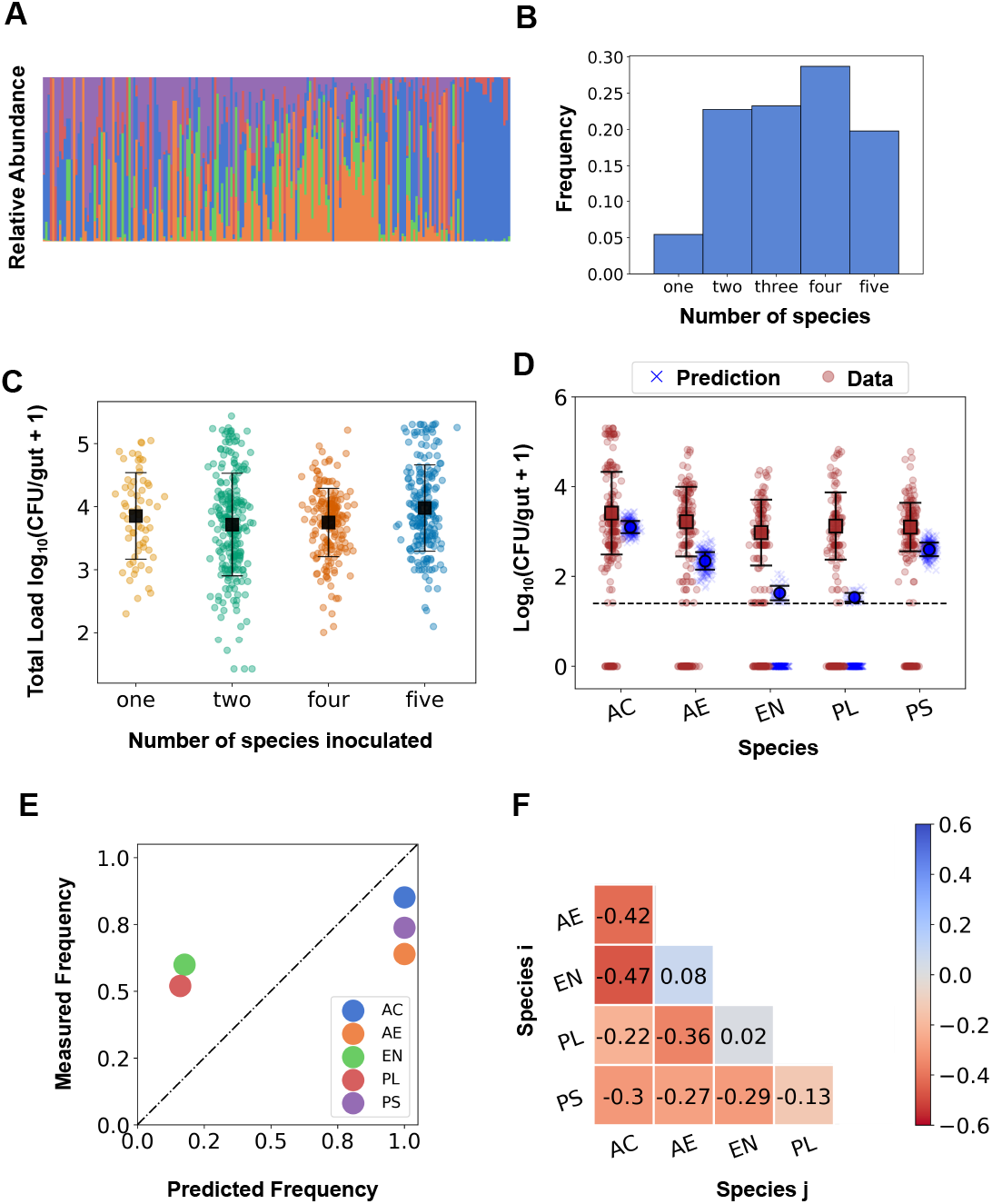
Communities are more diverse and abundant in five-species experiments than would be predicted solely based on two-species pairwise interactions. (A) Stacked bar plot of the relative abundances of the five bacterial species when all five were co-inoculated. Each bar is from a single dissected fish. The bars are ordered by total bacterial load. (B) Histogram of the total number of bacterial species present in the gut when all five species were co-inoculated. (C) The total bacterial load as a function of the number of inoculated species. Each circular datapoint is a CFU value from an individual fish (N = 63, 232, 187, and 202, from left to right), with the mean and standard deviation indicated by the square marker and error bars. (D) The predicted (blue Xs) and measured (brown circles) abundances of each bacterial species in the zebrafish gut when all five species are co-inoculated. Predictions are based on an interaction model that is linear in log-abundance using the pairwise 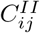 coefficients, as described in the main text. Solid square markers indicate the mean and standard deviation of the distributions excluding null counts. The dotted line indicates the experimental detection limit of 25 cells. The experimental data is from N = 202 fish in total and the predicted distributions arise from 250 samples of the distribution of interaction coefficients. (E) The observed frequency of occurrence in the gut from the five-species co-inoculation experiment versus the predicted frequencies for each of the five species. (F) The Pearson correlation coefficients calculated from the relative abundances of pairs of species when all five species were co-inoculated.

We plot the measured and predicted distributions of intestinal abundances of each of the five species for the five-species co-inoculation experiment in Fig 4D. The measured distributions of each of the species are very similar to each other. In contrast, the distributions of the predicted abundances vary significantly by species. For two of the species, AC and PS, the mean of the observed and predicted distributions are similar. For the other three species, in contrast, the observed and predicted populations are in strong disagreement, with the pairwise prediction being at least an order of magnitude lower than the observed abundances. For EN and PL in particular, we would expect extinction in a large fraction of fish due to strong negative pairwise interactions; in actuality, both species are common and abundant.

Similarly, we can extract from the model the predicted frequency of occurrence of each of the species, regardless of abundance. We find that the predicted frequency is much lower than the experimentally observed frequency for PL and EN (Fig 4E).

By measuring absolute abundances of bacterial populations in the gut, we provide direct assessments of inter-species interactions. More common sequencing-based methods, applied for example to the human gut microbiome, typically provide relative measures of species abundance, i.e. each taxonomic unit’s fraction of the total load. Correlations among relative abundances are often used as measures of interaction strengths [59, 60]. Calculating the Pearson correlation coefficients of the relative abundances of each pair of species in fish inoculated with all five bacterial species, we find a strikingly different interaction matrix (Fig 4F) than that inferred from absolute abundance changes (Fig 4B), with many strong negative values. By definition, the correlation matrix is symmetric (*C_ij_* = *C_ji_*), which does not capture the asymmetry inferred from absolute abundance data (Figs 2C, 3B).

## Discussion

Using a model system comprising five commensal bacterial species in the larval zebrafish intestine, we have characterized aspects of gut microbiome assembly. Controlled combinations of inoculated species and measurements of absolute abundance in the gut, both challenging to perform in other vertebrate systems, reveal clear signatures of interactions among species. We find strong, competitive interactions among certain pairs in fish inoculated with two bacterial species. In contrast, pairwise interactions are weak in intestines colonized by four to five species, and all species are present at equal or greater abundance than would be predicted based on two-species data.

Our quantification of interaction strengths relies on a minimal set of assumptions that serve as a general test of additive models. Interaction strengths are necessarily parameters of some model. In the text, we make use of a model in which the log-transformed population of a species is a linear function of the other species’ log-transformed populations, and a more general power law model that spans both sub-linear and super-linear dependences on population sizes. There are good reasons to be skeptical of such frameworks. First, intestinal populations may not be well described by equilibrium, steady-state values. Second, these models lack spatial structure information. In vivo microscopy of one or two species in the zebrafish gut [39, 40, 61] underscores both of these concerns: populations are very dynamic with rapid growth and stochastic expulsions; interactions can be mediated by complex intestinal mechanics; and aggregation and localization behaviors are species-specific.

Imaging also, however, provides justifications for these rough models. Prior microscopy-based studies have shown that growth rates are rapid, with populations reaching carrying capacities within roughly 12 hours [39, 61], well below the 48 hour assessment time considered here. Because of strong aggregation observed in nearly all bacterial species, most individual bacteria residing in the bulk of clusters will not directly interact with other species, leading to interactions that are sub-linear in population size, suggesting a logarithmic or *α* < 1 power-law functional form. Furthermore, stochastic dynamics can be mapped onto robust average properties for populations [39, 62]. It is therefore reasonable to make use of simple models, not as rigorous descriptions of the system but as approximations whose parameters characterize effective behaviors. We note that all these issues also affect more commonly used models, such as standard competitive Lotka-Volterra models that are linear in population sizes. These models are often applied to gut microbiome data and used to infer interaction parameters [26, 63, 64] despite a lack of information about their realism. The power law model of interactions provides the strongest indication of the generality of our conclusions. Over a range of interaction forms extending from highly sublinear (*α* = 0.1) to super-linear (*α* = 2.0), strong competitive interactions are damped when four or five species are present (Fig 3), suggestive of higher order interactions among intestinal bacteria.

The ecological potential for higher-order or indirect interactions, i.e. interactions that cannot be reduced to pairwise additive components but rather result from the activities of three or more species, to be important determinants of community structure has been appreciated for decades [41, 42, 47]. Identification of higher-order interactions among constituent species is important for accurate prediction of responses to ecological perturbations such as species invasion or extinction, as well as functions of multi-species communities, as such features will not be adequately forecast by examination of direct interactions in subsystems [41, 65].

Inferring and quantifying indirect interactions in natural ecosystems is, however, challenging, calling for subtle and model-dependent statistical tests [41, 42, 66]. Constructed or manipulated systems enable more straightforward assays in which particular species are introduced or removed amid a backdrop of others. Several such systems involving macroscopic organisms [67–71], as well as microorganisms [32, 50] have uncovered significant indirect interactions. However, some studies of microbial communities have found weak or negligible higher order interactions [47, 48], including one study examining combinations of species introduced to the *C. elegans* gut [31]. The complexity of interactions in a vertebrate gut has remained unclear, and correlation-based methods for inferring interactions from sequencing-based data have assumed that pairwise interactions suffice [59, 72, 73].

Our measurements using gnotobiotic larval zebrafish, a model vertebrate, show strong pairwise interactions when only two bacterial species are present in the intestine and weak pairwise interactions when four to five species are present, indicating strong higher-order interactions (Fig 3). In many cases, the effect is evident from the raw data itself. For example, EN is strongly suppressed by AE if the two are inoculated together (Fig 2B). Comparing EN abundance in fish colonized by all species except AE with its abundance in fish colonized by all species, however, shows little difference, indicating that the EN-AE interaction is strongly attenuated by the presence of the other bacterial groups (Fig 3A).

Two additional observations also imply the presence of strong higher-order interactions in our intestinal ecosystem. Considering fish colonized by all five bacterial species, the abundance of each species is at least as high (Fig 4D) and the diversity of bacterial species is higher than the values that would be predicted solely from direct interactions (Fig 4E).

Our finding of increased species diversity in a system of several gut bacterial species is consistent with recent theoretical studies that suggest, for a variety of reasons, that higher-order interactions are likely to stabilize communities and promote coexistence. The topic of multi-species coexistence has a long history in ecology. Especially since classic work by Robert May showing that a system comprising pairwise interacting constituents will, in general, be less stable as the number of species increases [74], explaining how complex communities can exist has been a theoretical challenge. There are many resolutions to this paradox, such as spatial heterogeneity, interactions across trophic levels, and temporal variation. However, even without such additional structure, incorporating higher order terms into general random competitive interaction models leads to widespread coexistence [43–45]. Such large-scale coexistence can also emerge naturally from contemporary resource competition models [46, 75], in which cross-feeding or metabolic tradeoffs necessarily involve multiple interacting species. Intriguingly, the abundance distributions of all five of our gut bacterial species, when inoculated together, are similar to one another. The average Shannon entropy of the five species community (H = 1.16 ± 0.24) also resembles that of a purely neutrally assembled community (H = 1.61), reminiscent of dynamics mimicking neutral assembly that emerge from multi-species dynamics driven by resource use constraints [46, 76].

Our findings imply that measurements of two-species interactions among microbial residents of the vertebrate gut are likely to be insufficient for predictions of community dynamics and composition. Moreover, they imply that inference from microbiome data of inter-species interactions, for example by fitting Lotka-Volterra-type models with pairwise interaction terms [26, 63, 64, 77] should not be thought of as representing fundamental pairwise interactions that would be manifested, for example, if the constituent species were isolated, but rather as effective interactions in a complex milieu.

Our measurements do not shed light on what mechanisms give rise to higher-order interactions in our system. Likely candidates include metabolic interactions among the species, interactions mediated by host activities such as immune responses, and modulation of spatial structure by coexisting species. Immune responses are sensitive to specific bacterial species [78] and to bacterial behaviors [79]. Regarding spatial structure in particular, in vivo imaging of these bacterial species in mono-association has shown robust aggregation behaviors that correlate with location in the gut [55] Given the physical constraints of the intestinal environment, we think that modification of spatial organization due to the presence of species with overlapping distributions is a likely mechanism for higher order interactions. Notably, both immune responses and spatial structure are amenable to live imaging in larval zebrafish [39, 40, 55]. Though the parameter space of transgenic hosts, fluorescent labels, and interaction timescales to explore in imaging studies is potentially very large, such future studies are likely to yield valuable insights into the mechanisms orchestrating the strong interactions observed here. Furthermore, examination of the roles of priority effects and other aspects of initial colonization, as well as stability of diverse communities with respect to invasion, may reveal potential routes for intentionally manipulating the vertebrate microbiome to engineer desired traits.

## Supporting information

**S1 File Absolute abundance data for all in vivo experiments** Spreadsheet containing absolute counts of bacterial cells observed for the each of the species for each fish inoculated in 1,2,4 and 5 experiments. Data from each experiment is in a separate sheet titled with experiment name eg. ‘2-species’. Within each sheet, separated columns represent the species combination inoculated for that particular experiment.

**S2 File Absolute bacterial cell counts for all in vitro experiments** Spreadsheet containing absolute counts of bacterial cells/mL for in vitro mono-association (sheet 1-species) and di-association (sheet 2-species) experiments in Lysogeny broth (LB) media.

**S3 File Pairwise interaction coefficients** 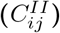 **for two species experiments calculated from the power law model** Spreadsheet containing the interaction coefficients for the two species experiments calculated using the power law model for different values of *α*. Each sheet contains the mean and standard deviation of the pairwise interaction coefficients for a single value of *α*. The first column in each sheet depicts the species pair (i-j).

**S4 File Pairwise interaction coefficients** 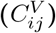 **for five species experiments calculated from the power law model** Spreadsheet containing the interaction coefficients for the five species experiments calculated using the power law model for different values of *α*. Each sheet contains the mean and standard deviation of the pairwise interaction coefficients for a single value of *α*. The first column in each sheet depicts the species pair (i-j).

**S5 File Predicted abundance distributions for log transformed abundance model** The file contains the predicted five species abundances of species using the linear additive model described in main text and the pairwise coefficients determined using the log transformed abundance model. Each row represents the predicted log_10_ abundance of species AC, AE, EN, PL and PS in one fish.

**S6 File Predicted abundance distributions for linear abundance model** The file contains the predicted five species abundances of species using the linear additive model described in S1 Text and the pairwise coefficients determined using the absolute abundance model. Each row represents the predicted absolute abundance of species AC, AE, EN, PL and PS in one fish.

**S1 Fig.**
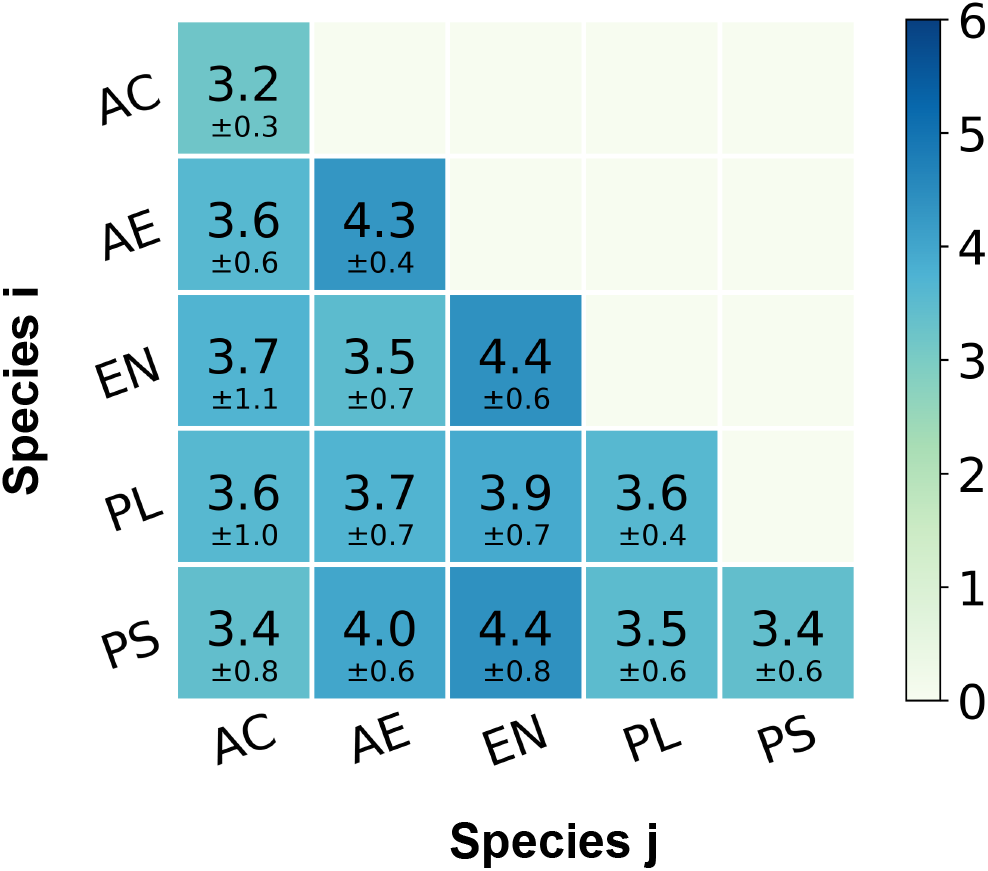
The total load for di-association experiments. The total bacterial load for different di-association experiments, expressed as the mean and standard deviation of log_10_ (total CFUs). The values on the diagonal are the mean load from mono-association experiments.

**S2 Fig.**
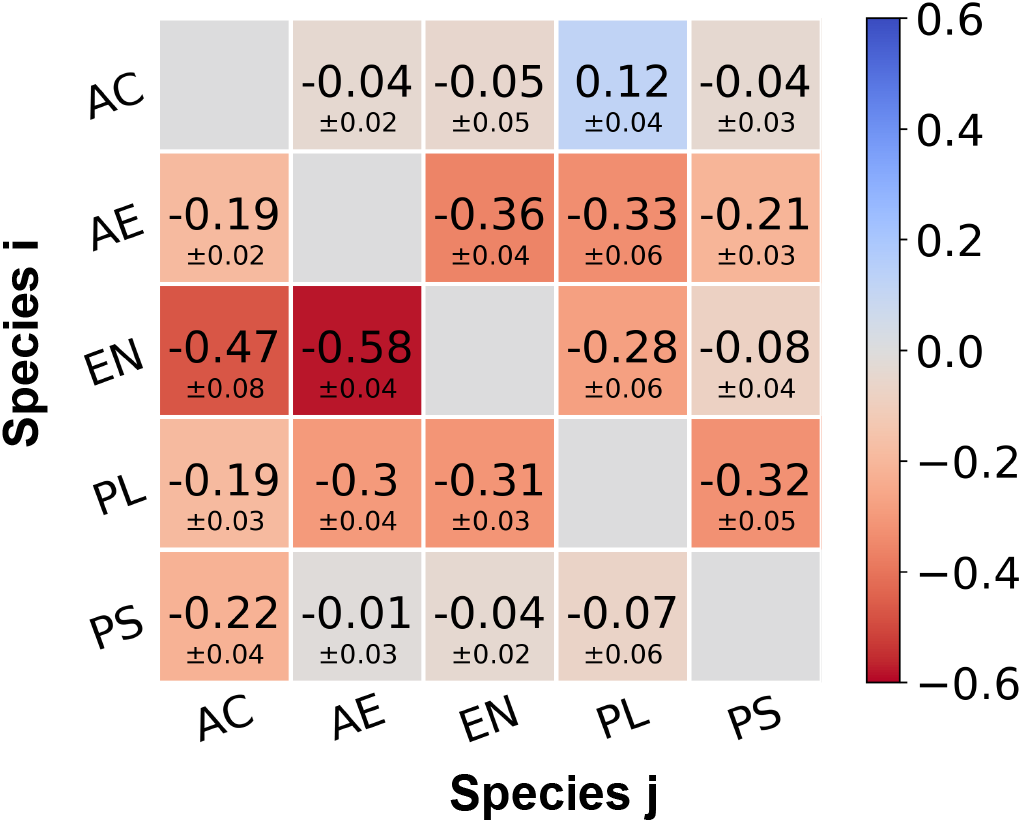
Pairwise interaction coefficients for two-species experiments using the log-transformed model. The mean pairwise interaction coefficients 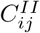 showing the effect of species j on species i calculated using the log-transformed abundance model. The standard deviations are calculated using a subsampling approach described in S1 Text.

**S3 Fig.**
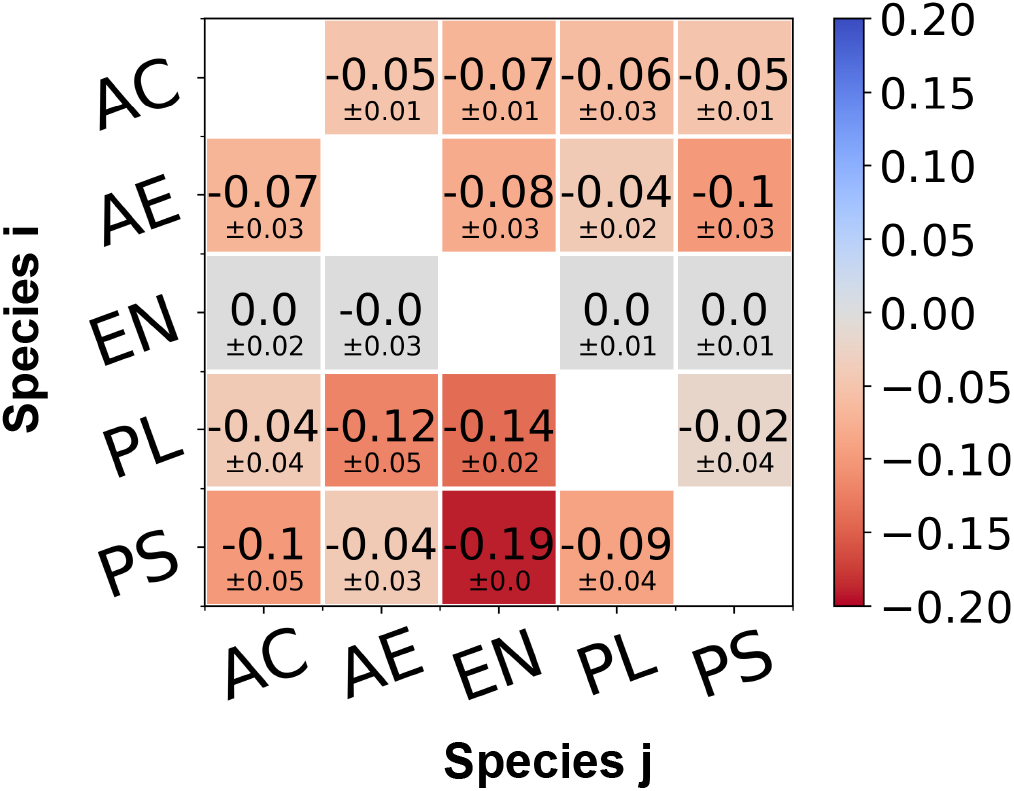
Pairwise interaction coefficients from in vitro two-species experiments. The matrix of interaction coefficients showing the mean and standard deviation of 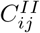 from in vitro competition experiments.

**S4 Fig.**
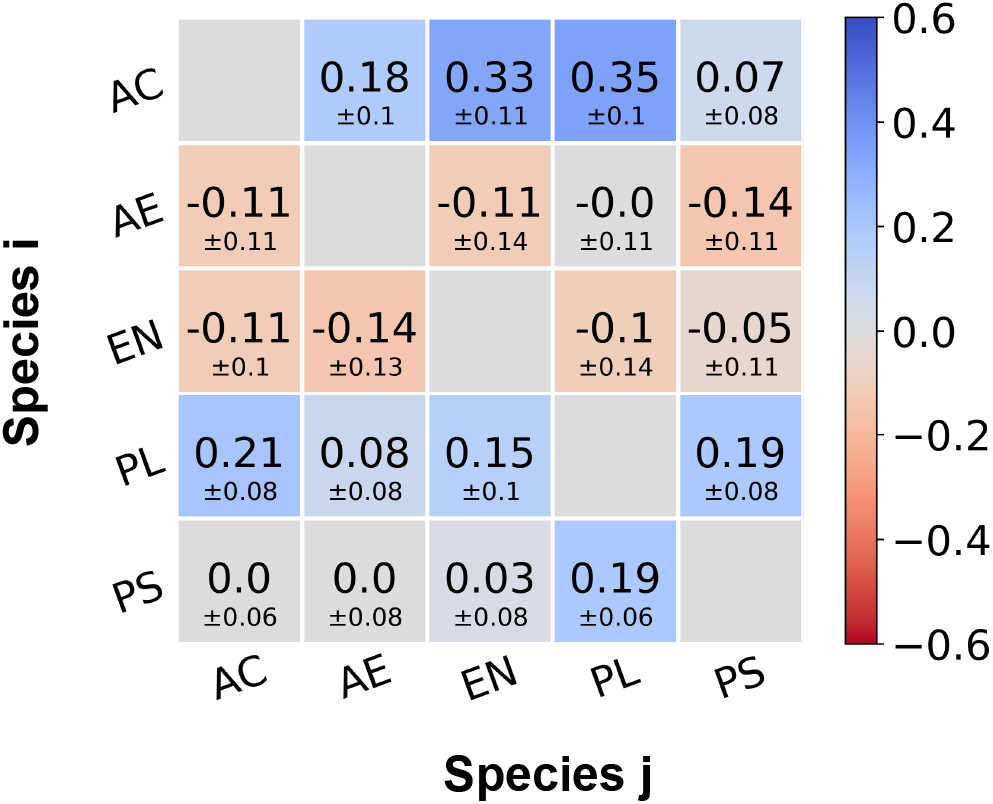
Pairwise interaction coefficients for five-species experiments using the log-transformed model. The mean pairwise interaction coefficients 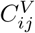 calculated using the log-transformed abundance model from experiments with four and five inoculated species. The standard deviations are calculated using a subsampling approach described in S1 Text.

**S5 Fig.**
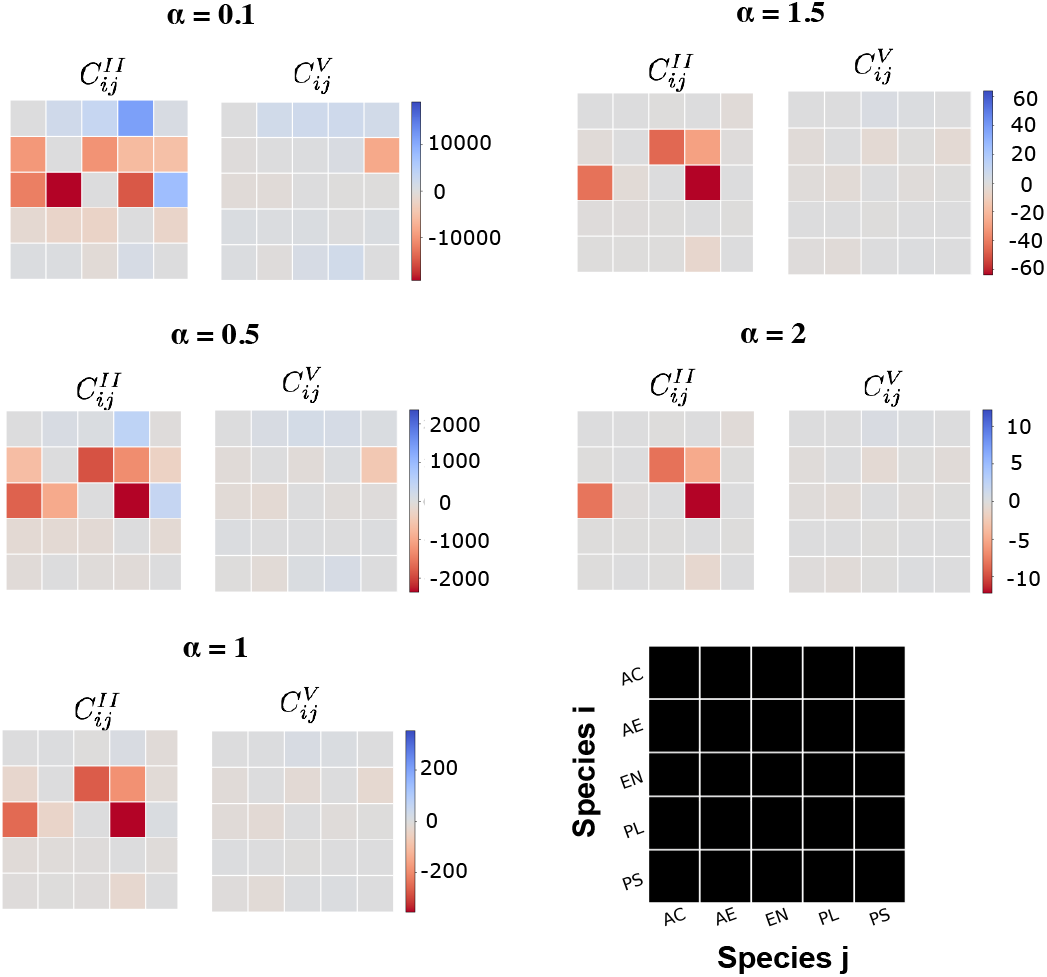
Pairwise interaction coefficients for select *α* values for two and five-species experiments using the power law model. Interaction coefficients generated from the linear absolute abundance model for are compared for the one-to-two species (left) and four-to-five species (right) experiments. The legend at the bottom-right shows the species labels for the rows and columns. The data is tabulated in S3 File and S4 File.

**S6 Fig.**
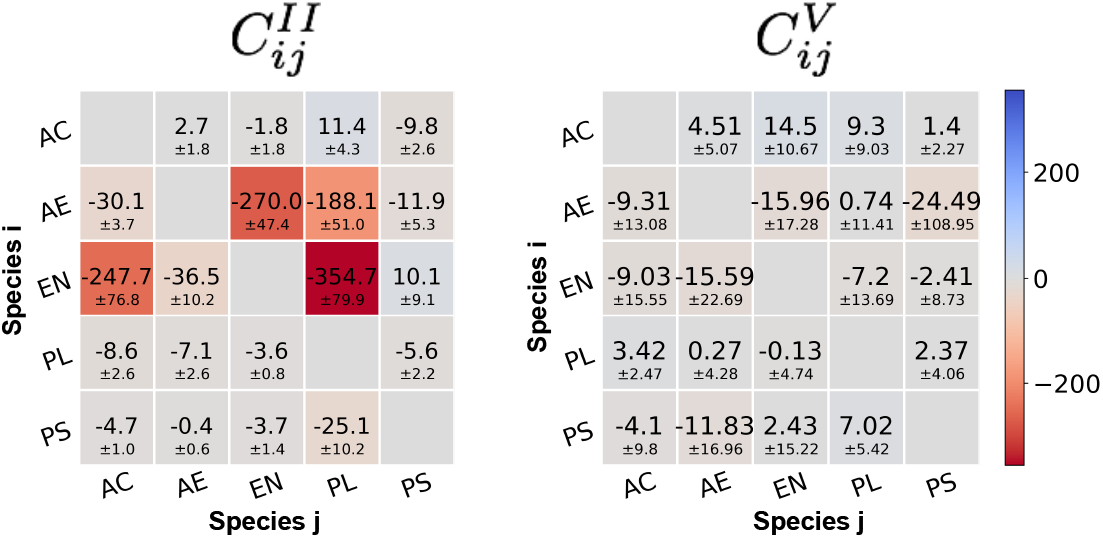
Pairwise interaction coefficients for two and five-species experiments using a linear model. The mean and standard deviation of the interaction coefficients for the two species 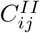 and five species 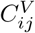 experiments calculated using a model linear in absolute species abundance.

**S7 Fig.**
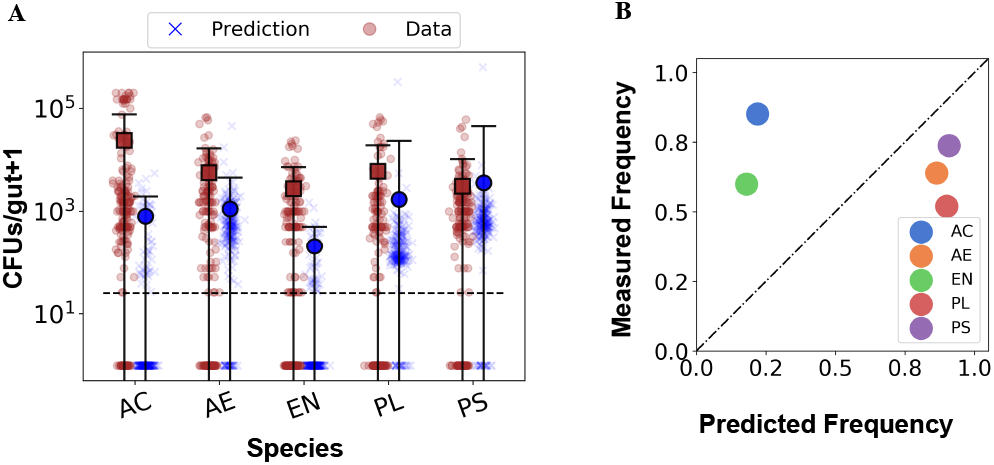
Predicted five species distributions from the linear model. A. Predicted (blue xs) and measured (brown circles) absolute abundance distributions for five-species inoculation experiments calculated from the linear model for pairwise interactions, for each of the five species. The means of the predicted and measured distributions excluding nulls are shown using bold blue circles and brown square markers, respectively, with error bars indicating the standard deviation. The dotted line indicates the experimental detection limit of 25 cells. The predicted distributions are generated from sampling the interaction coefficient distributions as described in S1 Text, while the experimental distributions comprise abundances from N = 202 fish. B. The observed occurrence frequencies of each species in five-species experiments plotted against the predicted frequencies generated from the linear model.

**S1 Text. Math Supplement** File containing a detailed description of the different interaction models and the procedures followed for carrying out the analysis of the experimental data.

## Acknowledgments

We thank Rose Sockol and the University of Oregon Zebrafish Facility staff for fish husbandry. We also thank Laura-Taggart Murphy, gnotobiology technician at University of Oregon for germ-free derivations of zebrafish larvae. Additionally, we also thank Jayson Paulose, Assistant Professor of Physics at the University of Oregon for insightful comments and suggestions.

