## Supplementary Text 1 for "Quantifying multi-species microbial interactions in the larval zebrafish gut"

### Math Supplement

#### 1 Interaction Models

##### 1.1 Pairwise Interactions

As noted in the main text, we aim to characterize the measured abundances of microbial species with numerical parameters that characterize their interactions. These parameters may simply be phenomenological, but may also relate to dynamical population models.

Considering two species, labeled by subscripts i and j, the commonly used competitive Lotka-Volterra model describes the behavior of the populations P by the following relationship:

$$\frac{\mathrm{d}P_i}{\mathrm{d}t} = r_i P_i \left(1 - \frac{P_i - C_{ij} P_j}{K_i}\right) \tag{1}$$

where $r_i$ is the intrinsic growth rate, $K_i$ is the carrying capacity, and $C_{ij}$, defined for is the interaction coefficient, negative for competitive interactions and positive for beneficial interactions.

As steady state, $\frac{\mathrm{d}P_i}{\mathrm{d}t} = 0$, hence,

$$C_{ij} = \frac{P_i - K_i}{P_j}. \tag{2}$$

Note that this holds even if the initial factor in the Lotka-Volterra equation, $r_i P_i$ is replaced by any generic function of $P_i$ . As discussed in the main text, each of the bacterial species examined is known to reach its carrying capacity approximately 10-12 hours after inoculation [1], a timescale considerably shorter than the 48 hours post-inoculation of our abundance measurements. It is therefore reasonable to make use of the steady-state approximation.

In our experiments, we measure intestinal abundances in di-associations, $P^{II}$, and mono-associations, $P^{I}$. The carrying capacity, $K_i$, is unknown. We assume $K_i$ can be well approximated by the average $P_i^I$ across fish, i.e. that the average abundance of a species in the absence of other species is similar to its carrying capacity in the gut. Therefore:

$$C_{ij} \approx \frac{P_i^{II} - \langle P_i^I \rangle}{P_j^{II}} \tag{3}$$

where the angle brackets indicate the mean across fish. Eq. 3 provides a simple characterization of inter-species interactions in the context of the standard competitive Lotka-Volterra equations.

As discussed in the main text, there is no reason the intestinal microbial populations must be governed by Lotka-Volterra dynamics or any linear interaction model. We therefore consider a model of logarithmic interactions and a more general family of power-law models. The logarithmic model, influenced by the order-of-magnitude changes in species abundance induced by competition, can follow from a phenomenological recasting of the competitive Lotka-Volterra model in terms of log-transformed populations:

$$\frac{\mathrm{d}\log_{10} P_i}{\mathrm{d}t} = f(P_i)\left(1 - \frac{\log_{10} P_i - C_{ij}\log_{10} P_j}{\log_{10} K_i}\right), \tag{4}$$

where we note that the first factor on the right hand side can be any generic function of $P_i$. Again assuming steady state dynamics and estimating $\log_{10} K_i$ by the average of $\log_{10} P_i$, the interaction coefficients are given by:

$$C_{ij} \approx \frac{\log_{10} P_i^{II} - \langle \log_{10} P_i^{I} \rangle}{\log_{10} P_j^{II}}. \tag{5}$$

which is Eq. 1 of the main text. We note that regardless of the validity of the dynamical model, Eq. 4 can be considered as a phenomenological characterization in which interaction effects are logarithmically additive (i.e. multiplicative) in population size.

Finally, we consider interactions with a power-law dependence on the size of the influencing species:

$$\frac{\mathrm{d}P_i}{\mathrm{d}t} = f(P_i)\left(1 - \frac{P_i - C_{ij}(P_j)^{\alpha}}{K_i}\right), \tag{6}$$

from which

$$C_{ij} \approx \frac{P_i^{II} - \langle P_i^{I} \rangle}{(P_j^{II})^{\alpha}} \tag{7}$$

As noted in the main text, we consider sub-linear ($\alpha < 1$) and super-linear ($\alpha > 1$) interactions. Linear interactions ($\alpha = 1$) are equivalent to the standard Lotka-Volterra model.

#### 1.2 Pairwise Interactions in the Presence of Additional Species

To compare interactions between communities of two species and communities of more species, namely four or five, we extended the above parameterization. The standard Lotka-Volterra dynamical model for an arbitrary number of species is given by

$$\frac{\mathrm{d}P_i}{\mathrm{d}t} = r_i P_i \left(1 - \frac{P_i - \sum\limits_{j\neq i} C_{ij} P_j}{K_i}\right) \tag{8}$$

where the sum runs over species. Again considering steady-state,

$$P_i = K_i + \sum_{j \neq i} C_{ij} P_j \tag{9}$$

As discussed in the main text, we infer $C_{ij}^V$ by comparing the abundance of species i in fish co-inoculated by a set of species excluding and including species j. Again denoting the number of species by a superscript, subtracting $P_i^{IV}$ from $P_i^V$ gives

$$P_i^V - P_i^{IV} = C_{ij}^V P_j^V, \tag{10}$$

from which

$$C_{ij}^V = \frac{P_i^V - P_i^{IV}}{P_j^V}. \tag{11}$$

Similar expressions follow for the multi-species interaction coefficient in the log-transformed model:

$$C_{ij}^V = \frac{\log_{10} P_i^V - \log_{10} P_i^{IV}}{\log_{10} P_j^V}. \tag{12}$$

and the power-law model:

$$C_{ij}^V = \frac{P_i^V - P_i^{IV}}{(P_j^V)^\alpha}. \tag{13}$$

The interaction matrices for calculated for the two-species experiments ($C_{ij}^{II}$), using Eq. 5 and four-to-five-species experiments ($C_{ij}^V$), using Eq. 12 are shown in S2 Fig and S4 Fig. As noted in the main text, we found similar patterns distinguishing $C_{ij}^{II}$ and $C_{ij}^V$, namely weaker and more positive interactions in the latter case, throughout the full range of $\alpha$ examined (Fig 3D, S5 Fig). The full distribution of mean $C_{ij}$ values for different values of $\alpha$ are provided in S3 File and S4 File.

### 2 Sampling and Parameter Estimation

For each di-association, we measured the species abundances $P_i^{II}$ and $P_j^{II}$ in N = 10 to 30 fish and calculated $C_{ij}^{II}$ for each according to Eq. 3, 5, 7 above. We did not consider data with $P_i^{II}$ or $P_j^{II}$ equal to zero to ensure that the measured abundance changes can be attributed to the presence of the second species. To determine the most likely $C_{ij}$ characterizing each inter-species interaction and its uncertainty, we randomly sampled 75% of each dataset without replacement and calculated the mean and standard deviation of the mean $C_{ij}$ from 3000 sampling repetitions. To determine the number of repetitions needed to accurately estimate $C_{ij}$, we performed the same analysis on simulated population data, drawing $P_i^{II}$ from a log-normal distribution and using a fixed $\langle \log_{10} P_i^I \rangle$. We then draw a value for $C_{ij}$ from a normal distribution of known mean and standard deviation and use this to compute $\log_{10} P_j^{II}$ using Eq. 5. Given the generated $\log_{10} P_j^{II}$ and $\log_{10} P_i^{II}$ distributions, we subsampled this dataset and computed the mean and standard deviation of the mean interaction coefficient varying the subsample size and number of repetitions. We found that on subsampling 75% of the dataset, $\sim 2000$ or more repetitions are needed to determine the standard deviation of $C_{ij}$ to an accuracy of 90%. The resulting mean and standard deviation of $C_{ij}^{II}$, one for each species pair, are shown in the figures in the main text and S2 Fig.

Interaction parameters in multi-species experiments, i.e. $C_{ij}^{V}$ were similarly calculated from subsampling abundance data from both the four species and the five species experiments. The mean and standard deviation of the $C_{ij}^{V}$ coefficients calculated using this model is shown in S4 Fig. As in the calculation of the di-association-derived $C_{ij}^{II}$, we considered in calculating $C_{ij}^{V}$ only data from fish in which each of the species inoculated in the four and five species experiments was detected, ensuring that the effects from all other species is consistent between experiments. In vitro interaction coefficients were also ascertained using Eq. 5 from abundance measurements in pairwise competition experiments in Lysogeny broth (LB) medium. The parameters shown in S3 Fig reflect the mean and standard deviation of the coefficients from six replicates for each of the pairs of species.

### 3 Relative Abundance Correlations

As noted in the main text, the Pearson correlation coefficients between the relative abundances of species are commonly interpreted as measures of interspecies interactions. Denoting the relative abundance of species i (i.e. the population normalized to the sum of all species populations) as $p_i$, we calculate the correlation coefficient as usual:

$$r_{ij} = \frac{\sum_{n=1}^{N} \Big((p_i)_n - \langle p_i \rangle\Big)\Big((p_j)_n - \langle p_j \rangle\Big)}{\sqrt{\sum_{n=1}^{N} \Big((p_i)_n - \langle p_i \rangle\Big)^2 \sum_{n=1}^{N} \Big((p_j)_n - \langle p_j \rangle\Big)^2}} \tag{14}$$

where N is the total number of fish examined. Note that by construction, $r_{ij}$ is symmetric.

### 4 Predicted Five-Species Abundance Distributions

We asked whether we can recreate the experimentally observed five-species abundance distributions using only the pairwise interaction coefficients from Section 1.1. Focusing first on the logarithmically additive model, we construct a predicted abundance value for each species by linearly combining the pairwise effects of each of the other species:

$$\log_{10} P_i^V = \langle \log_{10} P_i^I \rangle + \sum_{j \neq i} C_{ij}^{II} \log_{10} P_j^V . \tag{15}$$

$P_i^V$ and $P_j^V$ are the predicted abundances of species i and j respectively. This gives, for the five bacterial species in our system, five equations with five unknowns, namely the predicted $P_i$. To determine the distributions of predicted abundances for each of the five species, we randomly sampled each $C_{ij}^{II}$ from the interaction coefficient distributions obtained in Section 1.1 . We performed this sampling 250 times to generate a comparable number of points to the experimental distribution (N=202). Both the predicted and the measured distributions are shown in Fig 4C. The predicted abundance distributions are provided in S5 File. To mimic the experimental detection limit of approximately 25 cells, all predicted abundances below 25 were set to zero.

Similarly, we calculated predicted abundance distributions for using the linearly additive model in absolute abundance (i.e. competitive Lotka-Volterra), for which

$$P_i^V = \langle P_i^I \rangle + \sum_{j \neq i} C_{ij} P_j^V . \tag{16}$$

As above, we generated predicted distributions for each of the five species, shown in S7 Fig and the distributions are provided in S6 File.

As discussed in the main text, the predicted distributions from both pairwise-additive models indicate that the five species are less likely to coexist than is observed experimentally.
